# Dispensable regulation of brain development and myelination by the immune-related protein Serpina3n

**DOI:** 10.1101/2024.02.06.579239

**Authors:** Meina Zhu, Yan Wang, Joohyun Park, Annlin Titus, Fuzheng Guo

**Author notes:** Correspondence: Fuzheng Guo, UC Davis School of Medicine/IPRM Shriners hospital Sacramento CA. 2425 Stockton Blvd., Sacramento, CA 95817 ORCID: 0000-0003-3410- 8389.

## Abstract

Serine protease inhibitor clade A member 3n (Serpina3n) or its human orthologue SERPINA3 is a secretory immune-related molecule produced primarily in the liver and brain under homeostatic conditions and upregulated in response to system inflammation. Yet it remains elusive regarding its cellular identity and physiological significance in the development of the postnatal brain. Here, we reported that oligodendroglial lineage cells are the major cell population expressing Serpina3n protein in the postnatal murine CNS. Using loss-of-function genetic tools, we found that Serpina3n conditional knockout (cKO) from Olig2-expressing cells does not significantly affect cognitive and motor functions in mice. Serpina3n depletion does not appear to interfere with oligodendrocyte differentiation and developmental myelination nor affects the population of other glial cells and neurons *in vivo*. Together, these data suggest that the immune-related molecule Serpina3n plays a minor role, if any, in regulating neural cell development in the postnatal brain under homeostatic conditions. We found that Serpina3n is significantly upregulated in response to oxidative stress, and it potentiates oxidative injury and cell senescence of oligodendrocytes. Our data raise the interest in pursuing its functional significance in the CNS under disease/injury conditions.

## Introduction

Human SERPINA3 displays tissue-specific expression under homeostasis and may play diverse roles under physiological conditions (de Mezer et al., 2023; Zhu et al., 2024). Loss-of-function variants in SERPINA3 gene were reported to associate with cancer susceptibility (Koivuluoma et al., 2021) and generalized pustular psoriasis (Ortega et al., 2010) (Frey et al., 2020) in certain populations though the cause-and-effect relationship has not been established by animal studies. Furthermore, SERPINA3 is an established acute-phase protein, secreted primarily by hepatocytes, in response to systemic inflammation and has been proposed to regulate immune responses and systemic inflammation (Zhu *et al*., 2024). Murine Serpina3n was identified as the orthologue of human SERPINA3 based on structural and functional similarities (Horvath et al., 2005), open the possibility to use genetically modified Serpina3n mice to study its role in normal development and diseases. In the CNS, unbiased transcriptomic data suggest that *Serpina3n* mRNA is upregulated primarily in oligodendroglial lineage cells under various disease/injury conditions (Kenigsbuch et al., 2022) including normal aging and multiple sclerosis, an inflammatory demyelinating disorder in which oligodendrocytes and myelin are the primary targets for immune attacks.

Previous data reported that Serpina3n modulated CNS immune responses (neuroinflammation) presumably depending on its cellular origins: astrocyte-derived Serpina3n promoted whereas neuronal-derived Serpina3n diminished neuroinflammation and microglial activation (see review Zhu et a., 2014). It is now increasingly recognized that the developing brain is an organ of innate immune activation (Herrera-Rincon et al., 2020; Lenz and Nelson, 2018; Zengeler and Lukens, 2021) where the innate immune activity shapes proper brain development and animal behavior. Indeed, microglia, the brain resident immune cells, are constantly activated during normal brain development and plays crucial roles in supporting neuronal/synaptic development and oligodendroglial myelination (Cunningham et al., 2013; Nemes-Baran et al., 2020; Paolicelli et al., 2011; Ueno et al., 2013). Astrocytes, key players in regulating immune responses (Colombo and Farina, 2016), also modulate neuronal survival, synaptogenesis, and myelination via secreting various cytokines and growth factors during normal brain development (Kiray et al., 2016; Vivi and Di Benedetto, 2024). Thus microglia (astrocytes)-mediated innate immune activity is essential for brain development and animal behavior. Little is known about the role of the immune-related molecule Serpina3n in the normal developing brain.

Recently, SERPINA3/Serpina3n has been reported to express in embryonic radial glia cells and its overexpression impairs embryonic neurogenesis and cognitive function in mice (Zhao et al., 2022), highlighting its importance in embryonic brain development. It remains elusive 1) whether Serpina3n is expressed during postnatal CNS development and what cell types express Serpina3n, and 2) what the functional significance of Serpina3n is for postnatal brain development and animal behavior given its proposed regulatory role in immune responses.

We aimed to tackle these questions by assessing the developing brain of postnatal mice carrying Serpina3n knockout and wild-type alleles. We found that oligodendroglial lineage cells were the major cell type in the CNS white matter expressing Serpina3n protein in early developing brain. We employed *Cre-loxP* approach to deplete Serpina3n in Olig2-expressing cells (mainly oligodendroglial lineage cells) and assessed the effects of Serpina3n deficiency on oligodendroglial differentiation/myelination and the development of other glial/neuronal populations. Through our comprehensive assessments, we conclude that Serpina3n appears to play a minor role in normal brain development and animal motor/cognitive functions. Our study also provided a valuable genetic tool for future studies to decipher the pathophysiological role Serpina3n in CNS disease pathologies.

## Materials and Methods Transgenic mice

All mice were housed at 12 h light/dark cycle with free access to food and drink, and both males and females were used in this study. All transgenic mice were maintained on a C57BL/6 background and approved by Institutional Animal Care and Use Committee at the University of California, Davis. B6.129-Olig2tm1.1(cre)Wdr/J (Olig2-cre, RRID:IMSR_JAX:025567) (Schuller et al., 2008) and B6.129S-Serpina3ntm1.1Lbrl/J (Serpina3nfl/fl, RRID:IMSR_JAX:027511) mice (Vicuna et al., 2015) were purchased from JAX. Animal genotype was determined by PCR of genomic DNA extracted from tail tissue. All Cre lines were maintained as heterozygosity. All animals and procedures were approved by UC Davis IACUC (#23575 and #23752). All data measurement from correct genotypes were included in the data analysis without data exclusion. A total of 70 (please make sure it is correct) mice were used in the manuscript. The number of mice used in each figure panel was shown in the figure legends and summarized here. The number of mice used in each figure panel: Fig. 1A-B (6 Ctrl and 6 cKO), Fig. 1C-D (5 Ctrl and 4 cKO), Fig. 1E (17 Ctrl, 9 het-cKO and 24 cKO), Fig. 2 (8 Ctrl and 8 cKO), Fig. 3 (11 Ctrl, 10 het-cKO and 10 cKO), Fig. 4A-C (16 Ctrl, 10 het-cKO and 10 cKO), Fig. 4D-F (11 Ctrl, 10 het-cKO and 10 cKO), Fig. 5-6,8-9 (5 Ctrl and 5 cKO), Fig. 7 (3 Ctrl and 3 cKO), Fig. 10 (4 cKO and 5 Ctrl). (Meina, please correct the figure labelings since we changed a lot to the figures)

**Figure 1.**
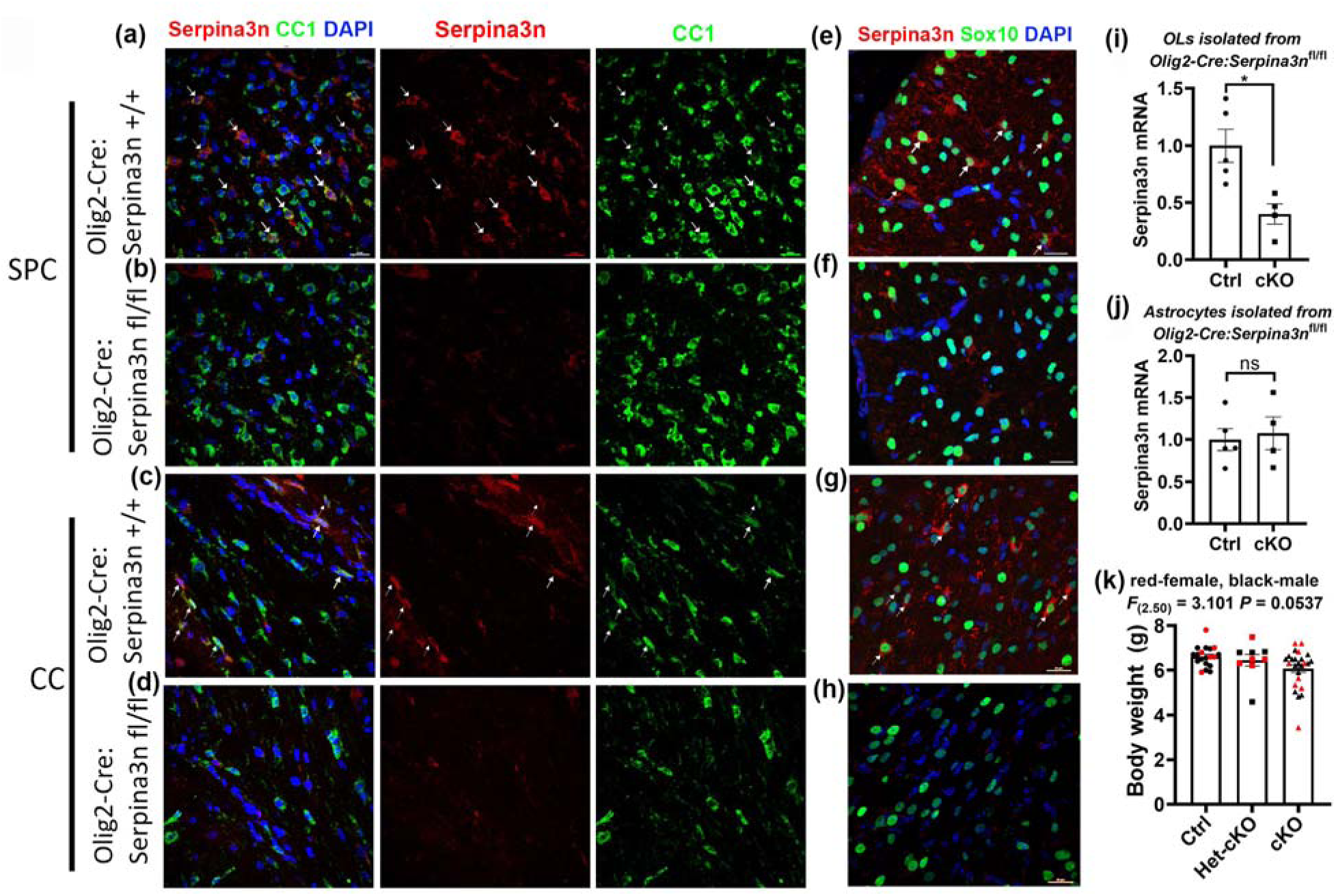
- Serpina3n expression and conditional knockout in the CNS. **(a, b, c, d)**, Serpina3n protein expression in CC1-positive OLs in the spinal cord (SPC, a, b) and corpus callosum (CC) (c, d) shown by fluorescent immunohistochemistry (IHC). Note that Serpina3n immunoreactive signal was eliminated in Serpina3n cKO mice. **(e, f, g, h)**, double IHC of Serpina3n and oligodendroglial lineage marker Sox10 in SPC (e, f) and CC (g, h). Note that almost all Serpina3n+ cells were positive for Serpina3n. Arrows in a-h pointing to double positive cells. **(i)**, RT-qPCR assay for Serpina3n in OLs acutely isolated by magnetic-assisted cell sorting (MACS, O4-magnetic beads) from P10 brain of Serpina3n cKO and Ctrl mice. Two- tailed Student’s t test, t_(7)_=3.285, * *P*<0.05. **(j)**, RT-qPCR assay for Serpina3n in astrocytes acutely isolated by MSCS (ACSA2-magnetic beads) from P10 brain of Serpina3n cKO and Ctrl mice. Two-tailed Student’s t test, t_(7)_=0.3281, *P*=0.7525. **(k)**, body weight of P14 mice. one-way ANOVA. *Olig2-Cre:Serpina3n*^fl/fl^ (Serpina3n cKO), *Olig2-Cre:Serpina3n*^fl/+^ (Serpina3n het-cKO) or *Olig2-Cre:Serpina3n*^+/+^ (Serpina3n Ctrl) mice were analyzed at postnatal day 10 (P10) in **panel A1-D** and P14 in **panel E.** Scale bars = 10 µm. (i, j) Ctrl N=5, cKO N=4; (k) Ctrl N=17, het-cKO N=9, cKO N=24.

**Figure 2.**
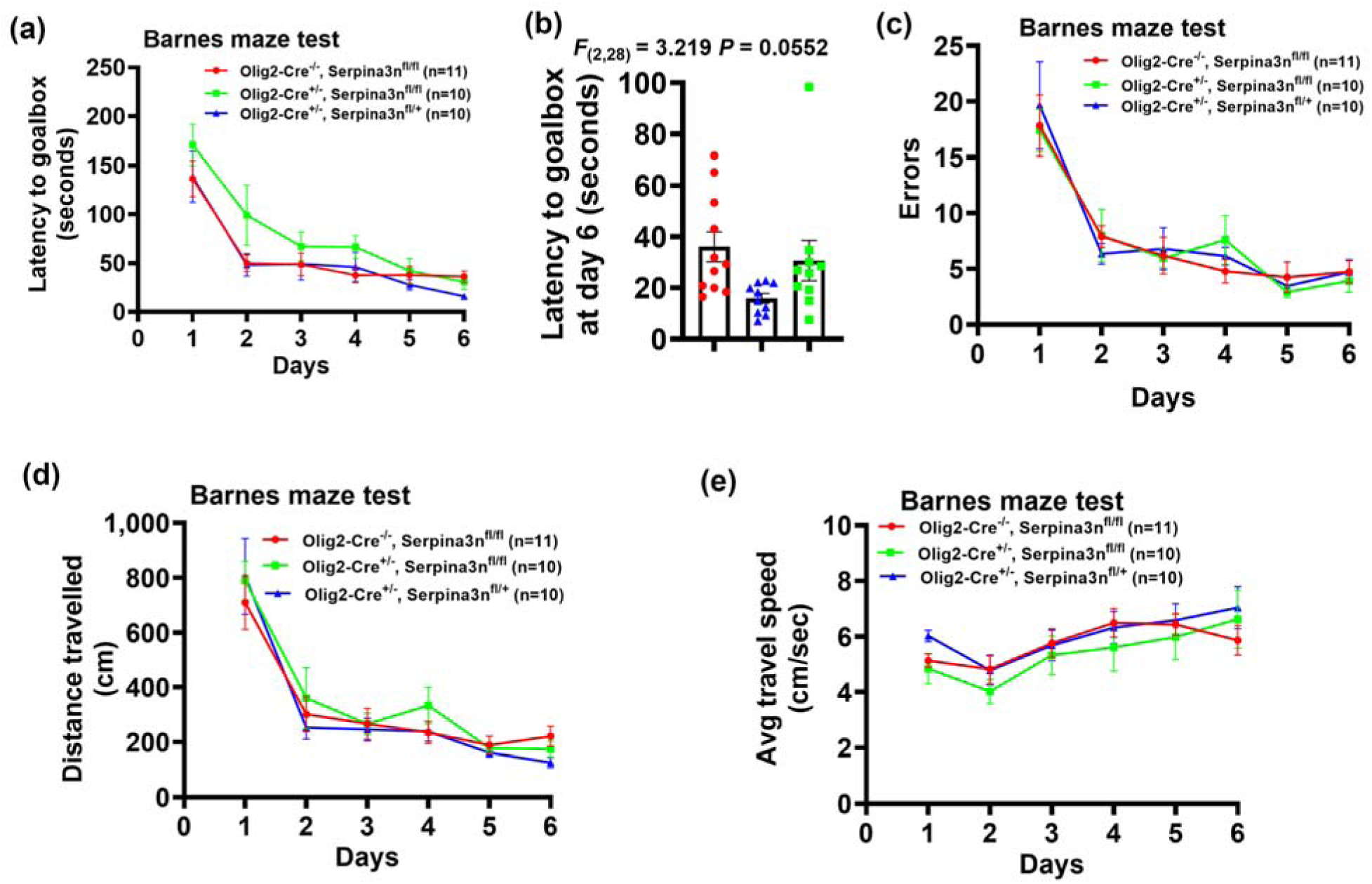
– Uninterrupted cognitive function of Serpina3n cKO mice. **(a)**, time latency (seconds) to find the goalbox. Time course *F*_(5, 168)_=27.37, *P*<0.0001, genotype *F*_(2, 168)_=6.2037, *P*=0.0025. * P<0.05 at Day 1. **(b)**, Latency to goalbox tested on day 6. **(c)**, number of errors made prior to goalbox. Time course *F*_(5, 168)_=29.38, *P*<0.0001, genotype *F*_(2, 168)_=0.037, *P*=0.9637. **(d)**, total distance travel during test. Time course *F*_(5, 168)_=38.50, *P*<0.0001, genotype *F*_(2, 168)_=0.7992, *P*=0.4523. **(e)**, average moving speed during test. Time course *F*_(5, 168)_=4.554, *P=*0.0006, genotype *F*_(2, 168)_=1.819, *P*=0.1654. Two-way ANOVA except B, one-way ANOVA.

**Figure 3.**
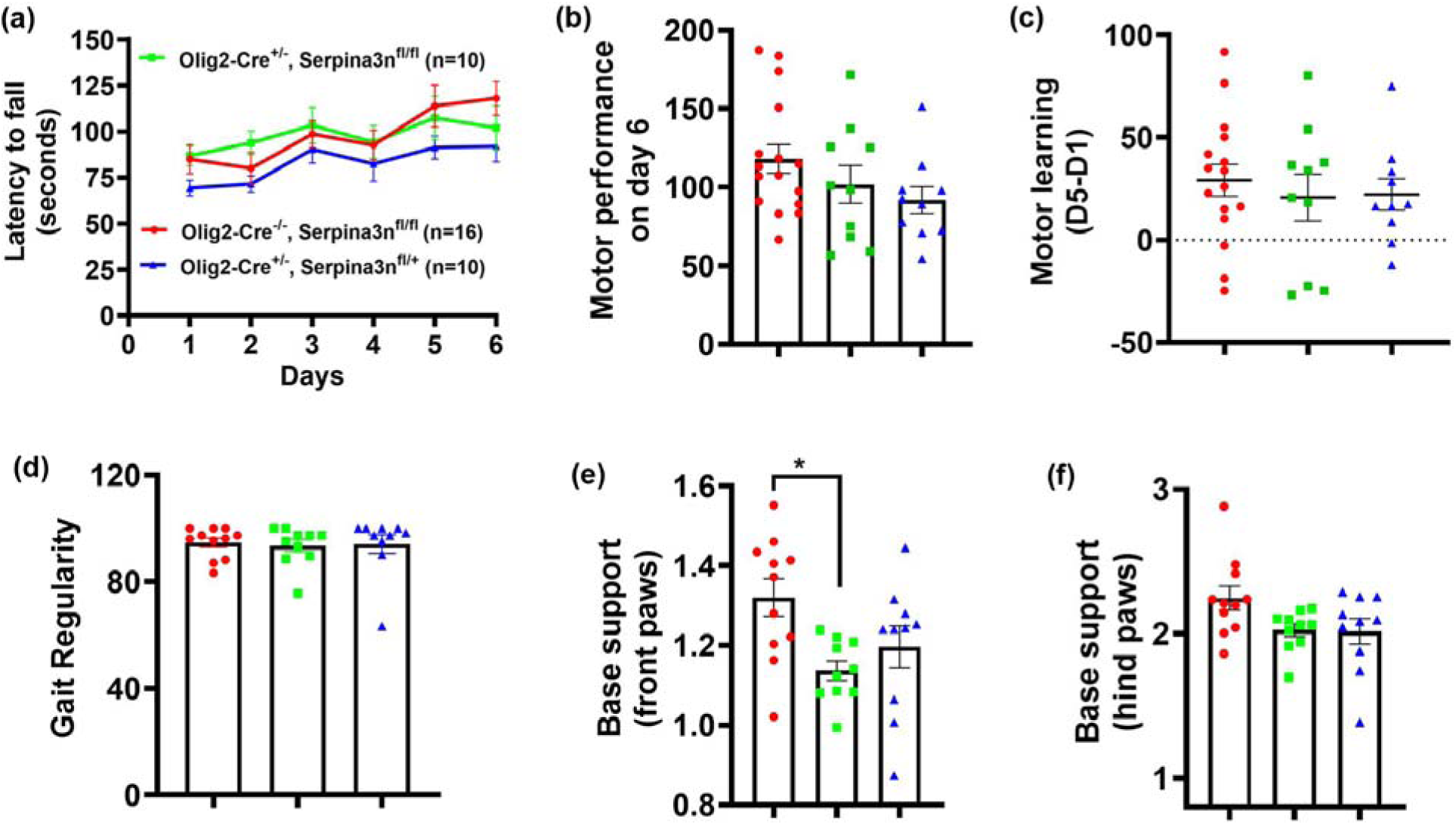
– motor function of Serpina3n-deficient mice **(a-c)**: motor function assessed accelerating Rorarod test; **(d-f)**: Gait analysis assessed by Noldus Catwalk. **(a)**, time-course of latency to fall (seconds) of mice during accelerating Rotarod test. Two-way ANOVA, time-course *F*_(3.4, 112.5)_=10.29, *P*<0.0001; genotype *F*_(2, 33)_=1.323, *P*=0.2801. **(b)**, motor performance on day 6 (latency to fall). One-way ANOVA, *F*_(2, 33)_=1.323, *P*=0.2801. **(c)**, motor learning ability (changes in latency to fall). One-way ANOVA, *F*_(2, 33)_=0.2811, *P*=0.7567. **(d)**, gait regularity index. One-way ANOVA, *F*_(2, 28)_=0.0531, *P*=0.9484. **(e-f)**, the base of support (the average width of two paws when walking) for front and hind paws (unit, centimeter). One-way ANOVA, front paws *F*_(2, 28)_=4.743, *P*=0.0168, hind paws *F*_(2, 28)_=3.115, *P*=0.0601, * P<0.05, Tukey’s multiple comparison test. Fig. 4a-c (Ctrl N=16, het-cKO N=10, cKO N=10), Fig. 4d-f (Ctrl N=11, het-cKO N=10, cKO N=10).

**Figure 4.**
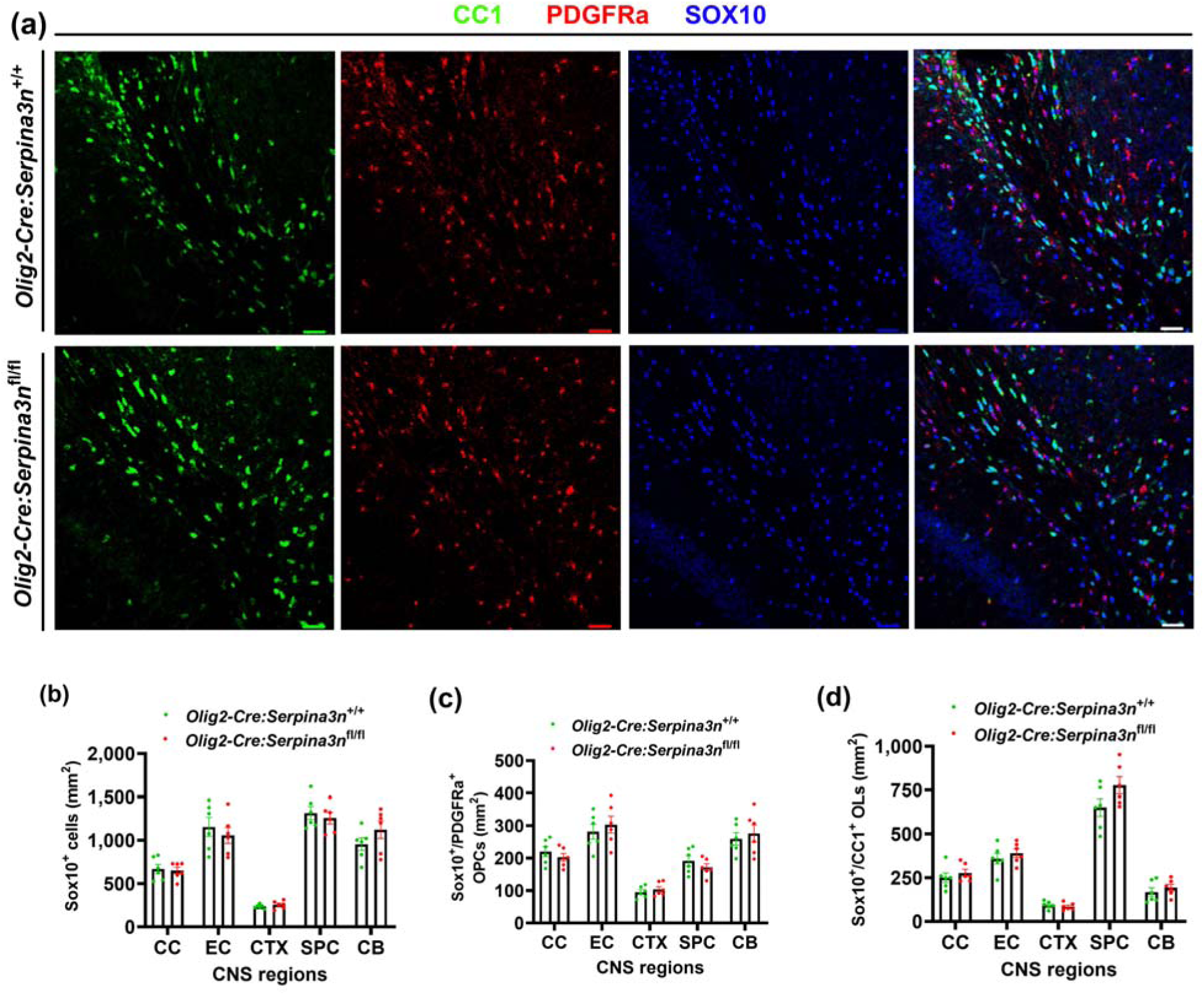
– oligodendrocyte differentiation during postnatal CNS development. **(a)**, representative confocal images pan-oligodendroglial lineage marker SOX10, differentiated OL marker CC1, and OPC marker PDGFRa in the corpus callosum (CC). **(b)**, quantification of pan-oligodendroglial lineage cells in different regions of the CNS. Two-tailed Student’s t test, CC t_(10)_=0.2806, P=0.2847; external capsule (EC) t_(10)_=0.6917, *P*=0.5048; cerebral cortex (CTX) t_(10)_=0.9386, *P*=0.3701; spinal cord (SPC) t_(10)_=0.5732, *P*=0.5792; cerebellum (CB) t_(10)_=1.354, *P*=0.2055. **(c)**, quantification of OPCs. Two-tailed Student’s t test, Sox10+/PDGFRa+: CC t_(10)_=8958, *P*=0.3914; EC t_(10)_=0.6342, *P*=0.5402; CTX t_(10)_=0.7610, *P*=0.4642; SPC t_(10)_=1.056, *P*=0.3156; CB t_(10)_=0.5225, *P*=0.6127. **(d)**, quantification of OLs. Two-tailed Student’s t test, Sox10+/CC1+: CC t_(10)_=0.4473, *P*=0.7909; EC t_(10)_=0.860, *P*=0.4390; CTX t_(10)_=0.7504, *P*=0.4703; SPC t_(10)_=1.8010, *P*=0.1019; CB t_(10)_=0.8010, *P*=0.4417. CNS tissues were harvested at P15. Scale bars = 50 µm. Ctrl N=6, cKO N=6.

**Figure 5.**
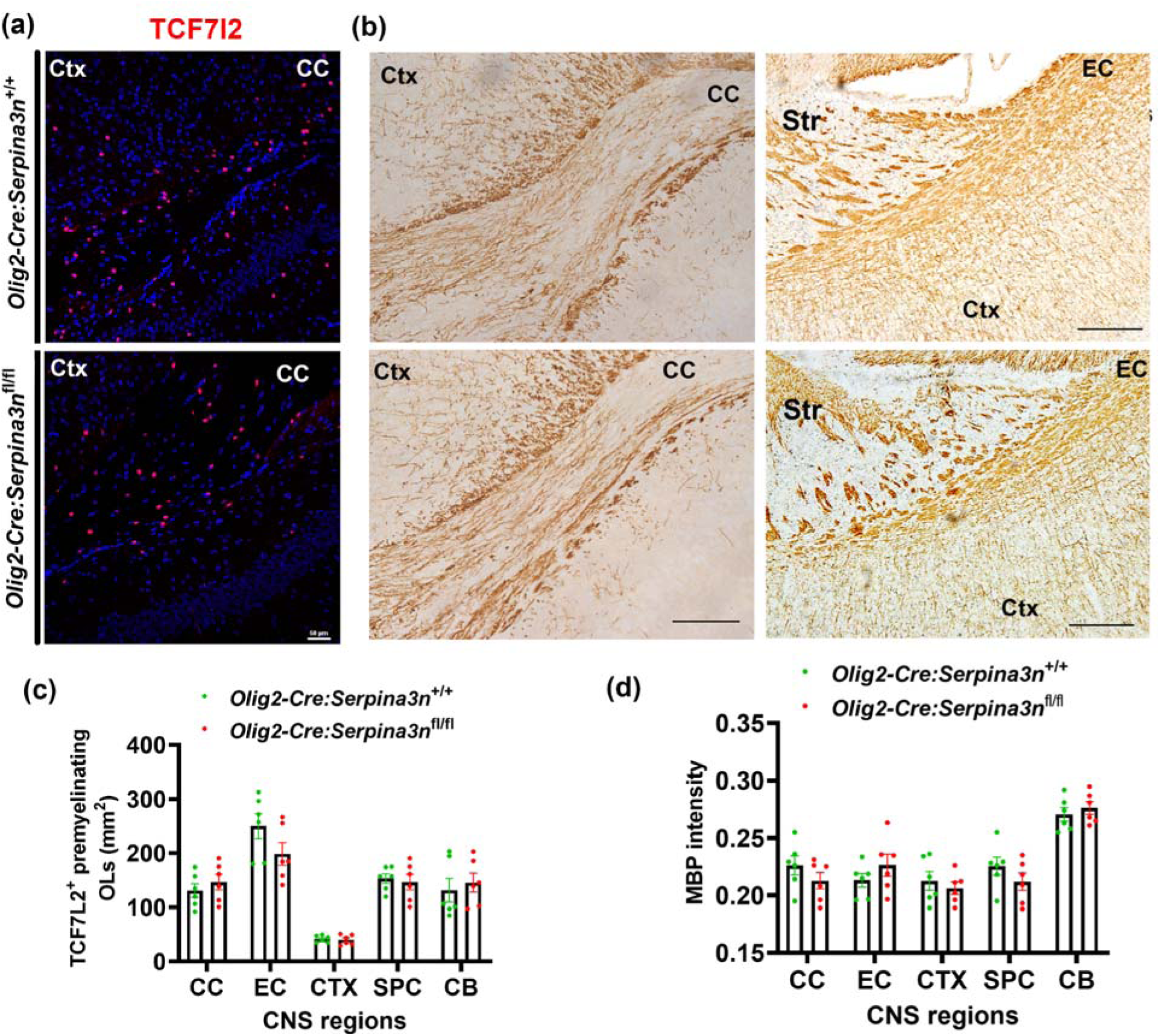
– Serpina3n deficiency does not affect the rate of oligodendrocyte genesis during normal development. **(a)** representative confocal images of newly generated premyelinating OL marker TCF7l2 in the CC. **(b)** DAB staining of myelin structure protein MBP in the CC and EC. **(c),** quantification of TCF7l2^+^ cells. Two-tailed Student’s t test, TCF4: CC t_(10)_=0.8234, *P*=0.4295; EC t_(10)_=1.646, *P*=0.1309, CTX t_(10)_=0.3702, *P*=0.7190; SPC: t_(10)_=0.3770, *P*=0.7141. CB t_(10)_=0.5059, *P*=0.6239. **(d)**, quantification of MBP intensity. Two-tailed Student’s t test, CC t_(10)_=1.220, *P*=0.2503; EC t_(10)_=1.170, *P*=0.2691; CTX t_(10)_=0.6571, *P*=0.5260; SPC t_(10)_=1.231, *P*=0.2466; CB t_(10)_=0.7021, *P*=0.4986. CNS tissues were harvested at P15. Scale bars = 50 µm. Ctrl N=6, cKO N=6.

**Figure 6.**
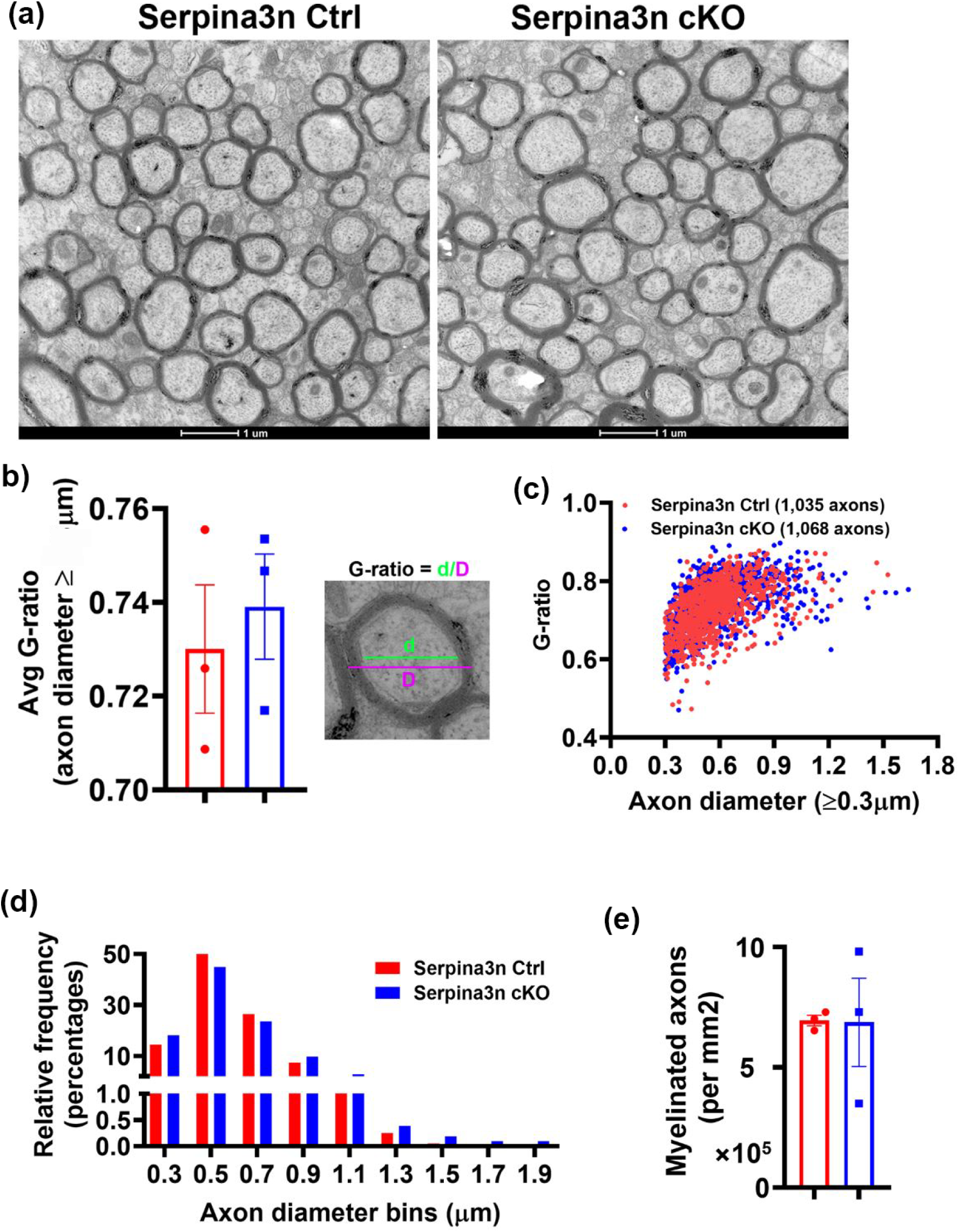
– normal myelination in Serpina3n cKO mice evaluated by transmission electron microscopy (TEM) **(a)**, representative TEM images from the corpus callosum demonstrating indistinguishable myelination profiles between Serpina3n Ctrl and cKO mice. Scale bars=1µm. **(b)**, Average g-ratio of myelinated axons in the CC. Two-tailed Student’s t test *t_(4)_*=0.5105, *P*=0.6366. **(c)**, scatter plot of G-ratio and axon diameters of individual axons (≥0.3µm in diameter). **(d)**, axon diameter frequency distribution. **(e)**, densities of myelinated axons. Two-tailed Student’s t test with Welch’s correction *t*_(2.059)_=0.0421, *P*=0.9702. The CNS tissues were harvested at P60. Ctrl N=3, cKO N=3.

**Figure 7.**
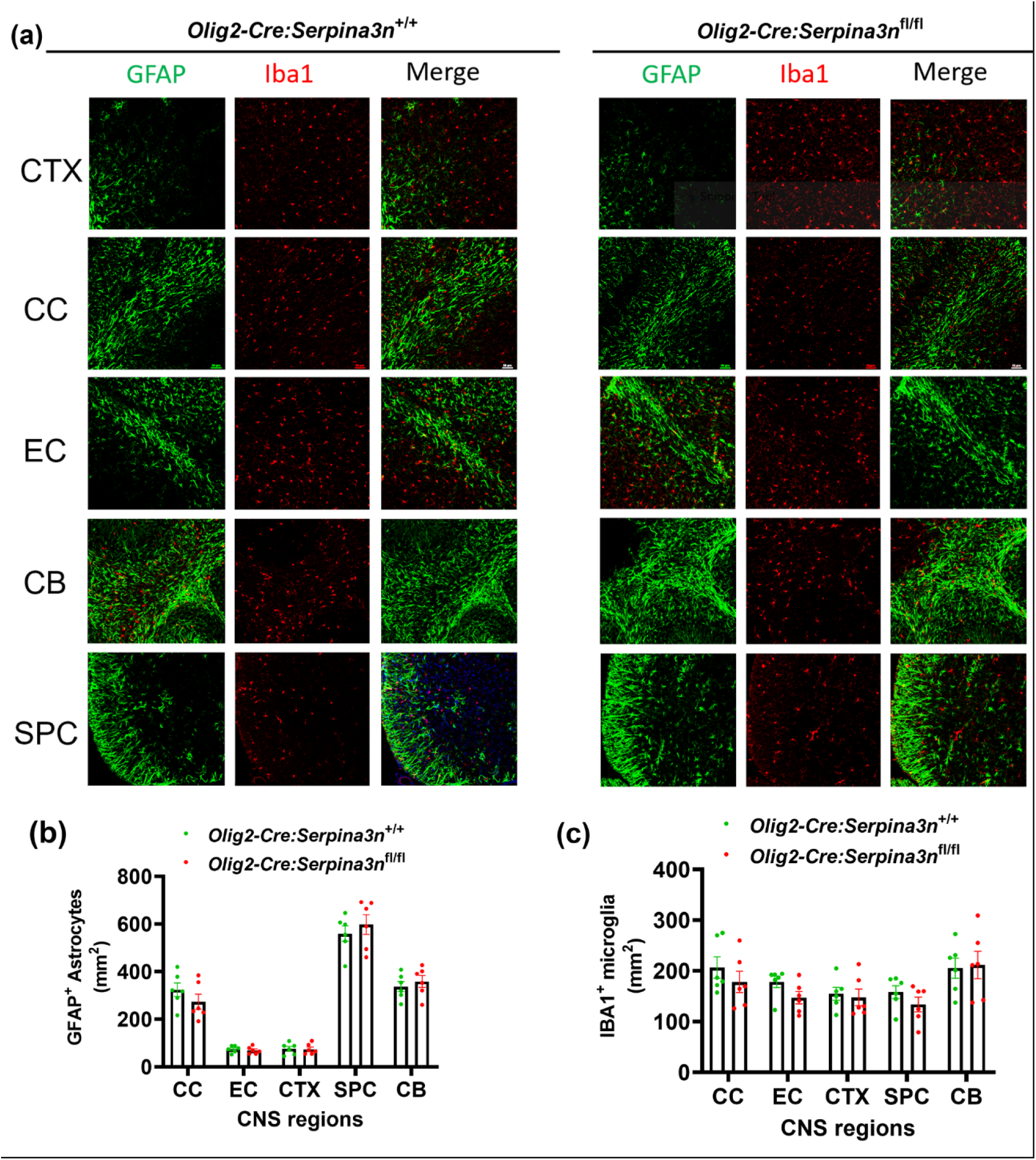
– Serpina3n deficiency does not affect astroglial or microglial development **(a),** representative confocal images of astroglial marker GFAP and microglial marker IBA1 in different CNS regions. **(b)**, quantification of GFAP+ astrocytes. Two-tailed Student’s t test, CC t_(10)_=1.177, *P*=0.2663; EC t_(10)_=0.6233, *P*=0.5740; CTX t_(10)_=0.1433, *P*=0.8889; SPC t_(10)_=0.7363, *P*=0.4785; CB t_(10)_=0.6508, *P*=0.0.5299. **(c)**, quantification of IBA1+ parenchymal microglia. Two-tailed Student’s t test, CC t_(10)_=0.9514, *P*=0.3638; EC t_(10)_=1.894, *P*=0.0875; CTX t_(10)_=0.3510, *P*=0.7329; SPC t_(10)_=1.271, *P*=0.2323; CB t_(10)_=1.271, *P*=0.2329. CNS tissues were harvested at P15. Ctrl N=6, cKO N=6.

## Tissue harvesting and processing

Serpina3n cKO mice and Ctrl mice were anesthetized by ketamine (150 mg/kg)/xylazine (15 mg/kg) mixture and transcardially perfused with ice-cold PBS. Harvested brain and spinal cord were placed immediately on dry ice for protein or RNA extraction or fixed in fresh 4% paraformaldehyde (PFA) in 0.1 m PBS overnight at 4°C for histological study. Then tissues were washed with PBS three times, 30 min per time and cryopreserved in 30% sucrose in PBS overnight at 4°C followed by embedding in O.C.T. (Cat# 361603E, VWR International). Serial sections (12 μm) were cut using a Leica Cryostat (CM1900-3-1). All slides were stored in −80°C freezers.

## Immunohistochemistry (IHC)

Sections were air-dried for 2 h and blocked in 10% donkey serum in 0.1% PBS Tween 20 (PBST) for 2 h at room temperature. For Serpina3n staining, slides were fixed with 100% methanol for 10 minutes at -20°C. Sections were washed with 0.1% PBST and incubated in primary antibody. After washing three times with PBST, slices were incubated with secondary antibodies for 2 h at room temperature. DAPI was used as a nuclear counterstain. Images were taken using a Nikon A1 confocal microscope. A z-stack of optical sections, 10 μm in total thickness, were collapsed into a single 2D image for quantification. The following antibodies were used for our IHC study: Goat anti-Serpina3n (1:3000, R&D System, Cat# AF4709, RRID:AB_2270116); Mouse anti-CC-1 (1:100, Millipore, Cat# OP80, RRID:AB_2057371); Goat anti-PDGFRα (1:200, R&D System, Cat# AF1062, RRID:AB_2236897); Rabbit anti-Sox10 (1:200, Abcam, Cat# ab155279, RRID:AB_2650603); Rabbit anti-TCF4/TCF7L2 (1:200, Cell Signaling Technology, Cat# 2569, RRID:AB_2199816); Mouse anti-MBP (1:200, MilliporeSigma, Cat# NE1019, RRID:AB_2140491); Mouse anti-GFAP (1:1000, Millipore, Cat# MAB360, RRID:AB_11212597); Rabbit anti-Iba1 (1:500, WAKO, Cat# 019-19741, RRID:AB_839504); Mouse anti-NeuN (1:500, Millipore, Cat# MAB377, RRID:AB_2298772). The signal was visualized by a secondary antibody Alexa Fluor 488- or Alexa Fluor 594-conjugated AffiniPure F(ab’)2 fragments (1:400, Jackson ImmunoResearch). Meina, include a brief description of DAB staining protocol and reagents

## Primary rodent oligodendrocyte culture

Primary oligodendrocyte cultures were established following our previously outlined protocols (#23575, 23752) (Zhang et al., 2021b). Eight pups per group at postnatal day 2 (P2) were anesthetized by hypothermia on wet ice (approximately 5–8 minutes in ice) followed by decapitation. The brains were dissected in ice-cold HBSS (#24020117, Thermo Fisher) under a microscope. Isolated tissues were pooled and underwent dissociation using a papain dissociation kit (#LK003176, Worthington) supplemented with DNase I (250 U/ml; #D5025, Sigma) and D-(+)-glucose (0.36%; #0188 AMRESCO) at in 37L℃/5% CO2 for 60 min. Tissue chunks were then transferred to the PDS Kit-Inhibitor solution (#LK003182, Worthington) and filtered through strainers. After centrifugation, the cell suspension was transferred to a poly-D-lysine (PDL, #A003-E, Millipore) pre-coated T-75 flask and maintained in DMEM high glucose (#11965118, Gibco) containing heat-inactivated fetal bovine serum (#12306-C, Sigma) and 1% penicillin/streptomycin (P/S, #15140122, Gibco) at in 37L℃/5% CO2 . The medium was replaced 50% every other day. When cells reached confluency around 11-14 days, the shaking method was applied to detach cells. To collect OPCs, microglia were detached with 220 rpm shaking for 2 hours at 37 ℃, and the supernatant was removed. 20 ml of serum-free growth medium (GM), a 3:7 mixture (v/v) of B104 neuroblastoma-conditioned medium, 10 ng/ml biotin (#B4639, Sigma), and N1 medium (high-glucose DMEM supplemented with 5 μg/ml insulin (#I6634, Sigma), 50 μg/ml apo-transferrin (#T2036, Sigma), 100 μM putrescine (#P5780, Sigma), 30 nM Sodium selenite (#S5261, Sigma), 20 nM progesterone (#P0130, Sigma) was added to each flask and shaken at 220 rpm for 6 hours. The supernatant was then collected into a 50 ml tube and centrifuged at 1800 rpm for 5 minutes. OPCs were cultured on PDL-coated plates with complete GM, consisting of GM with 5 ng/ml FGF (#450-33, Peprotech), 4 ng/ml PDGF-AA (#315-17, Peprotech), 50 µM forskolin (#6652995, Peprotech), and glutamax (#35050, Thermo Fisher). To induce differentiation, the medium was switched to differentiation medium (DM), composed of 12.5 μg/ml Insulin, 100 μM Putrescine, 24 nM Sodium selenite, 10 nM Progesterone, 10 ng/ml Biotin, 50 μg/ml Transferrin (#T8158, Sigma), 30 ng/ml 3,3′,5-Triiodo-L-thyronine (#T5516, Sigma), 40 ng/ml L-Thyroxine (#T0397, Sigma-Aldrich), glutamax, and P/S in F12/high-glucose DMEM, 1:1 in medium (#11330032, Thermo Fisher Scientific). After 3 days of differentiation, OLs were treated with PBS or 50 mM or 200 mM H_2_O_2_.

## Enzyme-linked immune sorbent assay (ELISA) of Serpina3n

The medium collected from primary culture were used for Serpina3n measurement assay at day 3 of differentiation. The Serpina3n concentration was determined using the mouse Serpina3n ELISA kit (BIOMATIK, EKF58884). ELISA was performed according to the manufacturer’s instruction.

## Magnetic-activated cell sorting (MACS)

Single-cell suspensions were prepared following the instructions provided by Miltenyi Biotec. Mouse brains from Control and Serpina3n cKO mice were processed using the Neural Tissue Dissociation Kit (P) (Cat# 130-092-628, Miltenyi Biotec, Germany) in conjunction with the gentleMACS Dissociator (Cat# 130-093-235, Miltenyi Biotec, Germany). The mouse brains were collected, cut into 0.5 cm pieces using a scalpel, and transferred into a pre-heated gentleMACS C tube (Cat# 130-093-237, Miltenyi Biotec, Germany). Enzymatic cell dissociation was initiated using 1950 µL of Enzyme mix 1 (Enzyme P and Buffer X). The C tube was attached upside down onto the sleeve of the gentleMACS Dissociator, and the brain tissue was dissociated using the appropriate gentleMACS program. After one rotation, 30 µL of enzyme mix 2 (Enzyme A and Buffer Y) was added into the C Tube, followed by two gentle rotations at 37°C. Upon program completion, the C Tube was detached, briefly centrifuged, and the sample at the bottom of the tube was filtered through a MACS SmartStrainer (70 μm) to remove cell clumps, achieving a single-cell suspension. The MACS SmartStrainer was washed with an additional 10 mL of HBSS (w) (HBSS with Ca2+ and Mg2+, Cat# 55021C, Sigma-Aldrich) to collect all the cells. After centrifugation, the supernatant was gently removed, and the brain homogenate pellet was incubated separately with Anti-CD11b Microbeads (130-093-634, Miltenyi Biotec, Germany) and Anti-O4 Microbeads (130-094-543, Miltenyi Biotec, Germany), with Fc receptors blocking prior to Anti-ACSA-2 MicroBeads (Cat# 130-097-678, Miltenyi Biotec, Germany). Incubation was carried out for 15 minutes in the refrigerator at 4℃. Cells were washed with 0.5% BSA/PBS buffer and centrifuged at 300 g for 10 min. The supernatant was aspirated completely, and the cell pellet was resuspended in 500 µL of 0.5% BSA/PBS buffer before proceeding to magnetic separation.

The LS MACS column was positioned in the magnetic field of a MACS Separator (Cat#130-090- 976, Miltenyi Biotec) and rinsed with 3 mL of 0.5% BSA/PBS buffer. The cell suspension was applied to the MS column, and the column was washed 3 times with 3 mL of 0.5% BSA/PBS buffer. Unlabeled cells were collected and combined with the flow-through. Magnetically labeled cells were immediately flushed out by firmly pushing the plunger into the column. Subsequently, 350 µL of Buffer RLT Plus containing β-mercaptoethanol (β-ME) was added to the cells, and the mixture was stored in a −80°C refrigerator.

## RNA extraction, cDNA preparation, and RT-qPCR

RNA isolation was carried out using the RNeasy Plus Micro Kit (Cat#74034, QIAGEN) following the manufacturer’s protocol. Harvested cells were transferred to a 1.5 mL tube and vortexed for 20 seconds. The lysate was then transferred to a gDNA Eliminator spin column placed in a collection tube. After centrifugation, 1 volume of 70% ethanol was added to the flow-through, mixed well by pipetting, and immediately transferred to RNeasy MinElute spin column placed in a collection tube. The RNeasy MinElute spin column was washed with Buffer RW1, Buffer RPE, and 80% ethanol, and the collection tube with the flow-through was discarded. RNase-free water was added to the spin column membrane and centrifuged for 2 minutes at 14,000 rpm to elute the RNA. The RNA concentration and purity (260/280 and 260/230 ratios) were analyzed using a Nanodrop 2000 Spectrophotometer (Thermo Fisher Scientific).

For cDNA synthesis, 200 ng of total RNA was reverse-transcribed using the QIAGEN Omniscript RT Kit (Cat#205111, QIAGEN) according to the manufacturer’s guidelines. The cDNA amplification was performed with a QuantiTect SYBR Green PCR Kit (Cat#204145, QIAGEN). Real-time PCR reactions were carried out and analyzed using an Agilent MP3005P thermocycler. For quantification, the mRNA expression level of target genes in each sample was normalized to that of Hsp90 as a reference gene. The fold change in gene expression level was calculated based on the equation 2^(Ct [cycle threshold][Hsp90] - Ct [target gene]). Primer sets were obtained from Integrated DNA Technologies.

## RT-qPCR primers

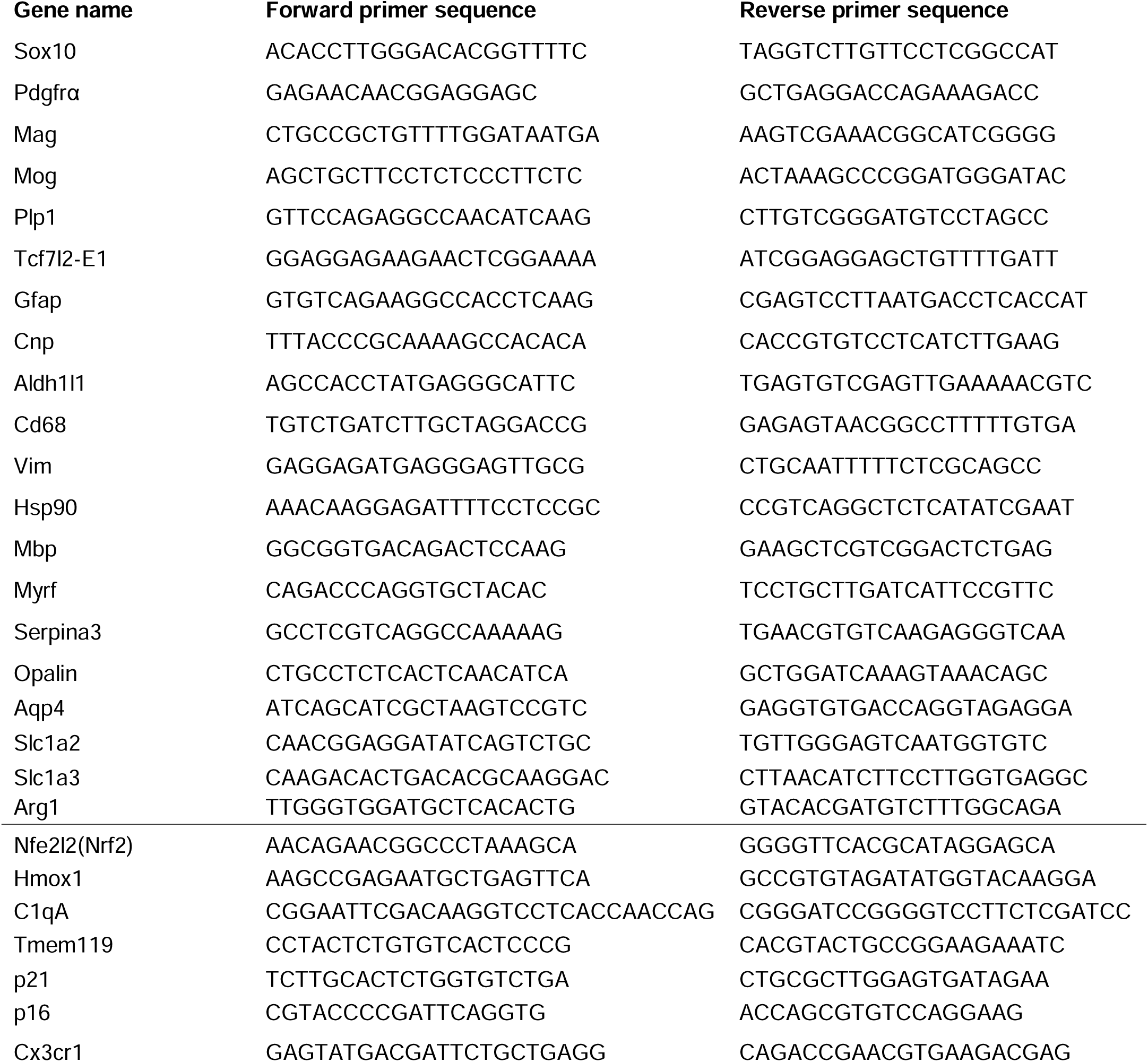
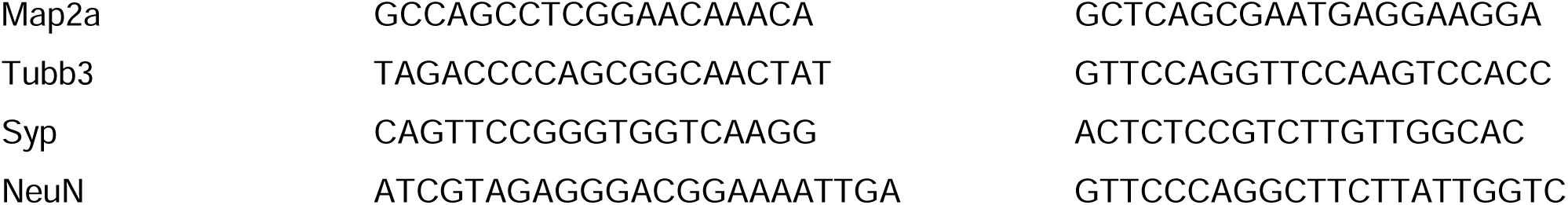

### Protein extraction and Western blot

Frozen tissues were homogenized in ice-cold N-PER Neuronal Protein Extraction Reagent (Thermo Fisher, #87792) containing protease inhibitor and phosphatase inhibitor cocktail (Cat# PPC1010, Thermo Fisher) and PMSF (Cat# 8553, Cell Signaling Technology) for 30 min and centrifuged at 14,000 rpm for 10 min at 4°C. Supernatants were collected and protein concentrations were measured using BCA protein assay Kit (Cat# 23225, ThermoFisher Scientific). Equal amounts of protein (30 μg) from each sample were loaded into the AnykD SDS-PAGE gels (BIO-RAD, # 4569036) for electrophoresis. The proteins were transferred onto a 0.2 μm nitrocellulose membrane (Cat# 1704158, BIO-RAD) using Trans-blot Turbo Transfer system (Cat# 1704150, Bio-Rad). The membrane was then blocked in 5% BSA for 1 h at room temperature, and incubated with primary antibodies overnight at 4℃ with Mouse anti-β-actin (1:1000, Cell Signaling Technology, cat#3700),Goat anti-SerpinA3N (1:500,LR&D Systems, AF7409), Goat anti-Olig2 (1:500,L Novus Biologicals, Cat# AF2418). After three times of 10 min wash in PBST, membranes were incubated with horseradish peroxidase (HRP)-linked donkey anti-goat or donkey anti-mouse antibody. The membrane was imaged using Western Lightening Plus ECL (Cat# NEL 103001EA, PerkinElmer). The intensities of the bands were quantified by ImageJ software to analyzing the scanned grayscale value.

### Transmission electron microscopy (TEM) for assessment of myelination

Mice were anesthetized with ketamine/xylazine mixture and perfused with 4% PFA, followed by 60 mL 3% glutaraldehyde (Electron Microscopy Science, dilute in PBS, pH 7.4) at a speed of 5 mL per minute. The brain was carefully dissected and fixed with 3% glutaraldehyde overnight. Subsequently, the brain underwent 2 washes with 0.2 M sodium cacodylate buffer (pH 7.2, Electron Microscopy Science), each lasting 10 minutes, followed by post-fixation with 2% (w/v) aqueous osmium tetroxide (Electron Microscopy Science) for 2 hours. After two additional washes with sodium cacodylate (10 minutes each), the brain was dehydrated using a gradient of ethanol (50%, 70%, 90%, and 100%), followed by 3 washes with propylene oxide (30 minutes each). The specimen was then incubated overnight in a 1:1 mixture of Propylene Oxide:Eponate Resin (Electron Microscopy Science) and subsequently treated with a 1:3 mixture of Propylene Oxide:Eponate Resin for 10 hours, followed by an overnight incubation in 100% Eponate Resin. The resulting specimens were embedded in EMBed-812 Resin for 2 days at 65℃.

Semithin sections (500 nm) were cut using a Leica EM UC6 microtome and incubated with 2% toluidine blue (Cat#194541, Ted Pella Inc.) at 100℃ for 2 minutes. These sections were then imaged using an Olympus BX61 microscope. Ultrathin sections (70–80 nm) were cut on a Leica EM UC7 microtome, collected on 1 mm Formvar-coated copper slot grids, double-stained with uranyl acetate and lead citrate, and imaged on a CM120 electron microscope.

### Animal Behavior assessment

Animals were acclimated to the behavioral room for 30 minutes before the tests. 12-week-old Serpina3n cKO and control mice underwent motor skill assessments using the CatWalk and Rotarod, followed by the Barnes maze cognition test.

### Accelerating Rotarod Test

The accelerating Rotarod test assessed animal motor coordination and performance according to established protocols. The Rotarod initiated at 4 rpm and accelerated to a maximum speed of 40 rpm, with a 1.2 rpm increment every 10 seconds.

Mice underwent two consecutive days of training (four trials each day with a 60-minute interval between trials), followed by data collection on the subsequent day. Each trial had a maximum duration of 300 seconds, and the time spent on the rod and maximal falling speed were recorded and averaged over the four trials.

### Noldus CatWalk Gait Analysis

Gait analysis was performed using the real-time video-tracked CatWalk XT system (Noldus Information Technology) within a walkway measuring 68 x 29 x 52.5 inches. Each animal underwent 3 runs in a day as a test session. Data collection parameters included a camera gain set to 20 and a detection threshold of 0.1, as per the manufacturer’s instructions. Successful runs were defined as those occurring within 0.50 to 5.00 seconds, and the average of three successful runs was used for data analysis.

### Barnes Maze Test

Spatial learning/memory and locomotion were evaluated using the Barnes maze. Mice were placed in the center of the maze (100 cm diameter) with twenty holes (10 cm diameter), aiming to locate the escaping (goal) box. The tests were conducted in a noise-free room with strong illumination (>300 LUX) and visual cues. Training spanned five consecutive days, with formal testing on day 6. Each animal underwent two trials per day with an approximately 60-minute interval. The maximal trial duration was 5 minutes, unless the mice found the goal box. EthoVision XT.14 software monitored and recorded various parameters, including total errors before entering the escape box, latency to entering the goal box, total distance traveled (path length), and moving velocity on the maze.

### Quantification and statistics

Quantification was performed by observers blind to genotypes and treatments. Data graphing and statistical analyses were performed using GraphPad Prism (version 8.0, www.graphpad.com). Data were presented as mean ± SEM in this study. Scatter dot plots were used to quantify data throughout our manuscript. Each dot in the scatter dot plots represents one mouse or one independent experiment. We used the online tool (https://clincalc.com/stats/samplesize.aspx) to calculate sample size and do power analysis using our previously publishe data. Unpaired two-tailed Student’s t test was used for statistically analyzing two groups of data and degree of freedom (df) were presented as t(df) in figure legends. Comparisons between more than two groups were analyzed by One-way ANOVA followed by Tukey’s post-test. Shapiro-Wilk approach determined data normality. F test and Browne-Forsythe test were used to test variance equality of two groups and three or more groups, respectively. The p-value was defined as *p < 0.05, **p < 0.01, ***p < 0.001, ns, not significant p > 0.05.

## Results

### Serpina3n expression in oligodendroglial lineage cells

We first asked what cell population(s) expresses Serpina3n in the homeostatic postnatal CNS. Double fluorescence immunohistochemistry (IHC) demonstrated that Serpina3n protein was clearly present in cells labeled by the monoclonal antibody CC1, an established marker for OLs, in the spinal cord (**Fig. 1a**) and the corpus callosum (**Fig. 1c**). In the spinal white matter tract, approximately 50% of CC1^+^ cells displayed Serpina3n immunoreactive signals and almost all Serpina3n^+^ cells were CC1 positive. To authenticate Serpina3n immunoreactive signals, we generated Serpina3n cKO mice by crossing Serpina3n-floxed mice with the widely-used oligodendroglial cell targeting Olig2-Cre line. *Olig2-Cre:Serpina3n*^fl/fl^ was referred to as Serpina3n cKO mice whereas *Olig2-Cre:Serpina3n*^fl/+^, Olig2-Cre, or Serpina3n^fl/fl^ as Serpina3n control (Ctrl) mice. Our data demonstrated that Serpina3n immunoreactive signals were mostly abolished in CC1^+^ OLs of Serpina3n cKO mice (**Fig. 1b, d**). Oligodendroglial Serpina3n expression was confirmed by double IHC showing that almost all Serpina3n^+^ cells were positive for Sox10 (**Fig. 1e, g**), a pan-oligodendroglial lineage marker and that Sepina3n was abolished in Serpina3n cKO mice (**Fig. 1f, h**). To prove the cellular specificity of Sepina3n cKO, brain OLs and astrocytes were acutely isolated by magnetic-assisted cell sorting (MACS) (Zhang et al., 2021a) (also see Fig. 9). We observed a significant reduction of Serpina3n mRNA in acutely isolated OLs (**Fig. 1i**) but not in astrocytes (**Fig. 1j**) of Serpina3n cKO mice compared to Serpina3n Ctrl mice. We failed to detect Serpina3n immunoreactive signals in the adult CNS (data not shown) except for the neurons in the basal-medial nucleus of the hypothalamus (Zhu *et al*., 2024). These data demonstrate that oligodendrocytes are the major cell population expressing Serpina3n in the early postnatal CNS. These expression data provided technical rationale to use *Olig2-Cre:Serpina3n*^fl/fl^ hybrids for studying the physiological function of Serpina3n in postnatal brain development.

### Serpina3n-deficient mice develop comparable cognitive and motor function to control mice

A recent study reported that SERPINA3 global overexpression augmented cognitive function in mice (Zhao *et al*., 2022). To decipher the physiological role of Serpina3n in animal behavior, we assessed our Serpina3n cKO and Ctrl mice for cognition and motor functions. Serpina3n cKO mice were born in expected Mendelian ratios and fertile. No gross growth abnormalities were observed during postnatal development and adult ages. The body weight of Serpina3n cKO mice were slightly reduced compared to Ctrl mice early postnatal ages (P14), but the difference was not statistically significant (**Fig. 1k**).

We employed Barnes Maze test to assess animal spatial learning function (Wang et al., 2022). Both Serpina3n cKO and Ctrl mice displayed remarkable improvement yet indistinguishable spatial learning ability during the training and probe sessions (**Fig. 2a-c**). No difference was observed in total distance traveled (**Fig. 2d**) or average travel speed (**Fig. 2e**) during 5-minute sessions. Taken together, these data suggest that Serpina3n plays a minor role in animal cognitive function during postnatal development.

We next employed accelerating Rotarod test to evaluate motor function. No difference in motor performance was observed between Serpina3n cKO and Ctrl mice during the training (**Fig. 3a**) and test (**Fig. 3b**) sessions. Serpina3n cKO mice also displayed normal ability of motor learning compared to controls (**Fig. 3c**). Sensitive CatWalk test with automatic video-tracking was used to test animal walking gait (Wang *et al*., 2022). The overall walking gait pattern was unaffected by Serpina3n deficiency evidenced by comparable gait regularity index (**Fig. 3d**).

The base of support of two front paws appeared to be affected by Serpina3n deficiency (**Fig. 3e**). However, the impairment was not seen in the hind paws of Serpina3n cKO mice (**Fig. 3f**).

These data suggest that oligodendroglial Serpina3n seems to play a minor role, if any, in mouse motor behavior.

### Normal oligodendrocyte differentiation in Serpina3n-deficient mice

Serpina3n was primarily expressed in OLs (**Fig. 1**). We therefore evaluated the potential intrinsic role of Serpina3n on oligodendroglial development. Triple fluorescence IHC of SOX10, PDGFRa, and CC1 (**Fig. 4a**) was used to quantify different stages of oligodendroglial lineage progression. The densities of total oligodendroglial lineage cells (SOX10^+^), OPCs (SOX10^+^/PDGFRa^+^), and differentiated OLs (SOX10^+^/CC1^+^) was assessed from different CNS regions. We chose P15 for quantification because oligodendroglial differentiation peaks in the CNS during this time window. Our thorough quantification failed to reveal significant differences in the number of oligodendroglial lineage cells (**Fig. 4b**), OPCs (**Fig. 4c**) or differentiated OLs (**Fig. 4d**) in various CNS regions. These data indicate that Serpina3n plays a minor role, if any, in oligodendroglial differentiation throughout the CNS.

### Unperturbed rate of oligodendrocyte differentiation in Serpina3n-deficient mice

In addition to assessing the accumulative OL population (**Fig. 4**), we next evaluate the rate of oligodendroglial differentiation to inform any potential defects of oligodendroglial development. To this end, we used TCF7l2, a nuclear marker that labels newly differentiated OLs and downregulation in mature post-myelinated OLs, thus reflecting the rate of oligodendrogenesis at any given time points (Guo et al., 2023). The nuclear expression of TCF7l2 makes quantification much easier than cell surface markers (**Fig. 5a**). No marked or statistically significant difference was observed in the densities of TCF7L2^+^ cells among different CNS areas (**Fig. 5c**), suggesting that Serpina3n deficiency did not affect the rate of oligodendrogenesis in vivo. The expression of myelin basic protein (MBP), one of the major structural proteins in myelin sheath, appeared to be comparable in most CNS regions between Serpina3n cKO and Ctrl mice (**Fig. 5b d**), suggesting that Serpina3n deficiency does not perturb myelin formation. Taken together, Serpina3n plays a minor role, if any, in regulating oligodendroglial differentiation.

### Myelination appears to proceed normally in Serpina3n-deficient mice

We next employed transmission electron microscopy (TEM) (Yan et al., 2022) to assess potential myelination abnormalities at the ultrastructural level at early adult P60 when developmental myelination is completed (**Fig. 6a**). G-ratio, the ratio of the inner axon diameter versus the total fiber diameter (inner axon+outer myelin) (**Fig. 6b, right**) was statistically comparable in Serpina3n-deficient mice to that in Serpina3n-sufficient mice (**Fig. 6b left, Fig. 6c**), suggesting that myelin sheath thickness of myelinated axons was unaffected by Serpina3n depletion. Furthermore, Serpina3n depletion did not alter the normal dynamics of axonal diameters (**Fig. 6d**). The density of myelinated axons (≥0.3µm) was also comparable between Serpina3n-deficient and control mice (**Fig. 6e**). These data demonstrate that oligodendroglial Serpina3n is dispensable for developmental myelination.

### Serpina3n deficiency does not perturb the normal development of astroglia, microglia or neurons

We reasoned that oligodendroglia-derived Serpina3n may regulate the development of other glial cells or neurons in a paracrine manner given its secretory nature. To this end, we next evaluated the development of microglial/astroglial and neuronal populations. Fluorescence IHC was used to quantify the population of astrocytes (GFAP) and microglia (Iba1) throughout different CNS regions (**Fig. 7a**). Our systemic quantification failed to reveal any statistically significant differences in the number of GFAP^+^ astrocytes (**Fig. 7b**) or Iba1^+^ microglia (**Fig. 7c**).

We next evaluate neuronal development in Serpina3n-deficient mice. IHC of NeuN, which labels neuronal cell bodies, and MAP2a, which labels neuronal dendrites and cell bodies, was used to quantify neuron population development (**Fig. 8a**). Our data showed that the density of neurons in the cerebral cortex was unaffected by Serpina3n deficiency (**Fig. 8b**). A similar conclusion was drawn by MAP2a quantification (**Fig. 8c**). Taken together, these quantitative data suggest that oligodendroglia-derived Serpina3n is dispensable for normal development of other glial cells and neurons in the CNS.

**Figure 8.**
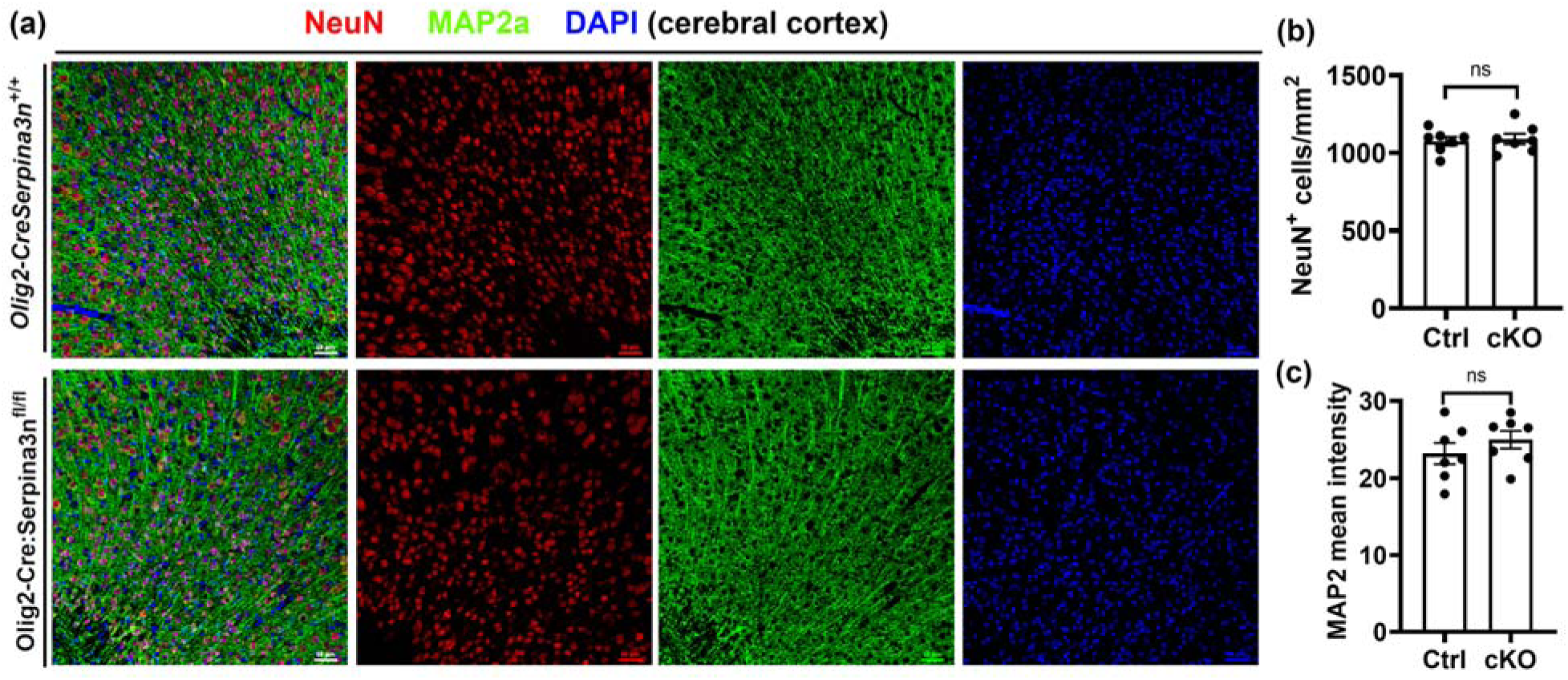
– Normal neuronal development in Serpina3n cKO mice. **(a),** representative confocal images of neuronal cell body marker NeuN and neurite marker MAP2a in the brain cerebral cortex. **(b-c)**, quantification of NeuN^+^ neuron population and MAP2a^+^ intentisy. Student’s t test, NeuN *t*_(12)_=0.3359, *P*=0.7427; Map2a *t*_(12)_=0.9995, *P*=0.3373. Brain collection at P15. Scale bars = 50 µm

### Molecular changes of brain cells in Serpina3n-deficient and control mice

We next aimed to quantify any potential alterations in glial cells and neurons at the molecular levels. For this purpose, magnetic-assisted cell soring (MACS) (Zhang *et al*., 2021a) was employed to acutely isolate microglia (CD11b), oligodendroglia (O4), astrocytes (ACSA2), and brain remnant cells (containing all neurons) (**Fig. 9a**). RT-qPCR assays of lineage-specific marker genes demonstrated that MACS-isolated cells were highly purified populations with little contamination of other cell types: *Tmem119* for microglia, *Sox10* for oligodendroglia, *Aldh1l1* for astroglia (**Fig. 19b**). RT-qPCR quantification of oligodendroglial mRNA showed no significant differences in the molecular expression of important functional genes indicative of oligodendroglial development, such as *Sox10, Pdgfra*, and *Tcf7l2* (**Fig. 9c**) and a panel of genes coding myelin proteins indicative of myelination (**Fig. 9d**). The expression levels of astroglial signature genes (*Gfap*, *Aldh1l1*) and crucial functional genes (*Slc1a2, Slc1a3, Aqp4*) were comparable between Serpina3n cKO mice and Ctrl mice (**Fig. 9e**). Moreover, microglial signature and functional genes were shown no significant differences between Serpina3n cKO mice and Ctrl mice (**Fig. 9f**). Because neurons were eluted into the triple negative cell component, we quantify neuronal marker genes (*Map2a, Tubb3, Syp, NeuN*) and found not significant difference in their mRNA expression between Serpina3n cKO and Ctrl mice (**Fig. 9g**). Altogether, these molecular assays demonstrate that Serpina3n plays a minor role, if any, in oligodendrocyte differentiation, myelination, and the development of other glial/neuronal populations.

**Figure 9.**
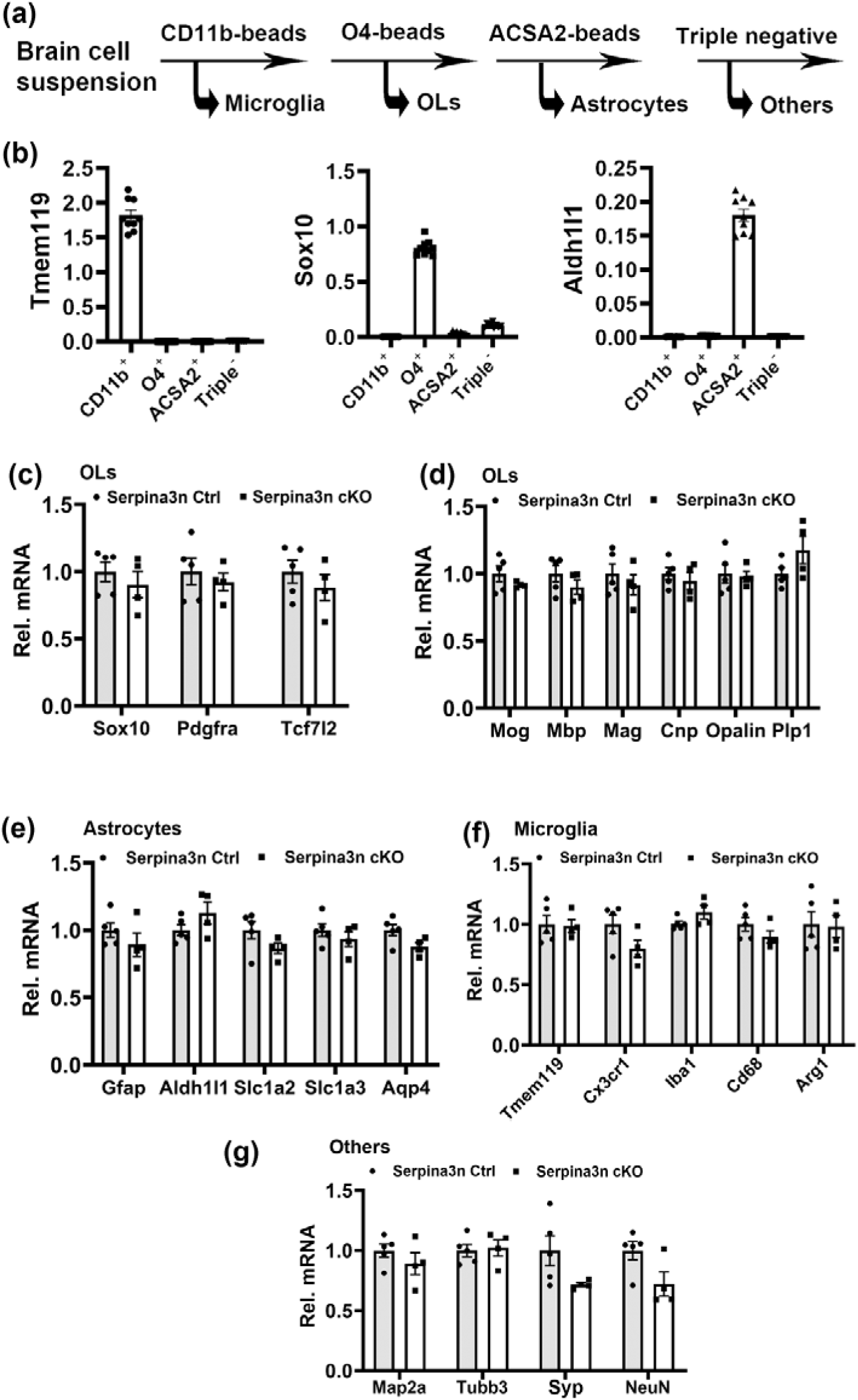
- unperturbed molecular expression of glial and neurons in acutely isolated cells **(a),** Experimental flowchart of acute isolation of CD11b^+^ microglia, O4^+^ OLs, ACSA2^+^ astrocytes, and triple-negative other brain cells by magnetic-assisted cell sorting (MACS) from P12 mouse brains. **(b)**, purity assays (by RT-qPCR) of each cell populations by lineage-specific markers, Tmem119 for microglia, Sox10 for OLs, and Aldh1l1 for astrocytes. **(c)**, RT-qPCR assay for the key genes of oligodendroglial development and differentiation in MACS-purified OLs. Sox10 *t*_(7)_=0.8067, *P*=0.4464; Pdgfra *t*_(7)_=0.6150, *P*=0.5580; Tcf7l2 *t*_(7)_=0.9178, *P*=0.3893. **(d)**, RT-qPCR assay for major oligodendroglial myelin-enriched genes in MACS-purified OLs. Mog *t*_(7)_=1.242, *P*=0.2542; Mbp *t*_(7)_=1.212, *P*=0.2645; Mag *t*_(7)_=0.7946, *P*=0.4529; Cnp *t*_(7)_=0.7261, *P*=0.4914; Opalin *t*_(7)_=0.2423, *P*=0.8155; Plp1 *t*_(7)_=1.721, *P*=0.1290. **(e)**, RT-qPCR assay for astrocyte marker genes in MACS-purified astrocytes. Gfap *t*_(7)_=1.102, *P*=0.3068; Aldh1l1 *t*_(7)_=1.509, *P*=0.1751; Slc1a2 *t*_(7)_=1.662, *P*=0.1405; Slc1a3 *t*_(7)_=0.9015, *P*=0.3973; Aqp4 *t*_(7)_=2.210, *P*=0.0628. **(f)**, RT-qPCR assay for microglial marker genes in MACS-purified microglia. Tmem119 *t*_(7)_=0.15352, *P*=0.8825; Cx3cr1 *t*_(7)_=1.914, *P*=0.0972; Iba1 *t*_(7)_=1.741, *P*=0.1251; Cd68 *t*_(7)_=1.352, *P*=0.2185; Arg1 *t*_(7)_=0.1326, *P*=0.8983. **(g)**, RT-qPCR assay for neuronal marker genes in triple-negative brain cells. Map2a *t*_(7)_=1.080, *P*=0.3160; Tubb3 *t*_(7)_=0.2726, *P*=0.7930; Syp *t*_(7)_=2.003, *P*=0.0553; NeuN *t*_(7)_=2.279, *P*=0.0567. Ctrl N=5, cKO N=4.

### Serpina3n is dispensable for oligodendrocyte differentiation *in vitro* but it potentiates oxidative stress and cell senescence under injury conditions

To determine the cell-autonomous effect of Serpina3n on oligodendrocyte differentiation, we employed primary oligodendrocyte culture. Serpina3n mRNA levels appeared to be increased in differentiating OLs (**Fig. 10a**) yet the protein level seemed to be unchanged during the lineage progression and maturation (**Fig. 10b**). We purified OPCs from neonatal brains of Serpina3n cKO and Ctrl mice and verified Serpina3n depletion in the cKO culture (**Fig. 10c**).

**Figure 10.**
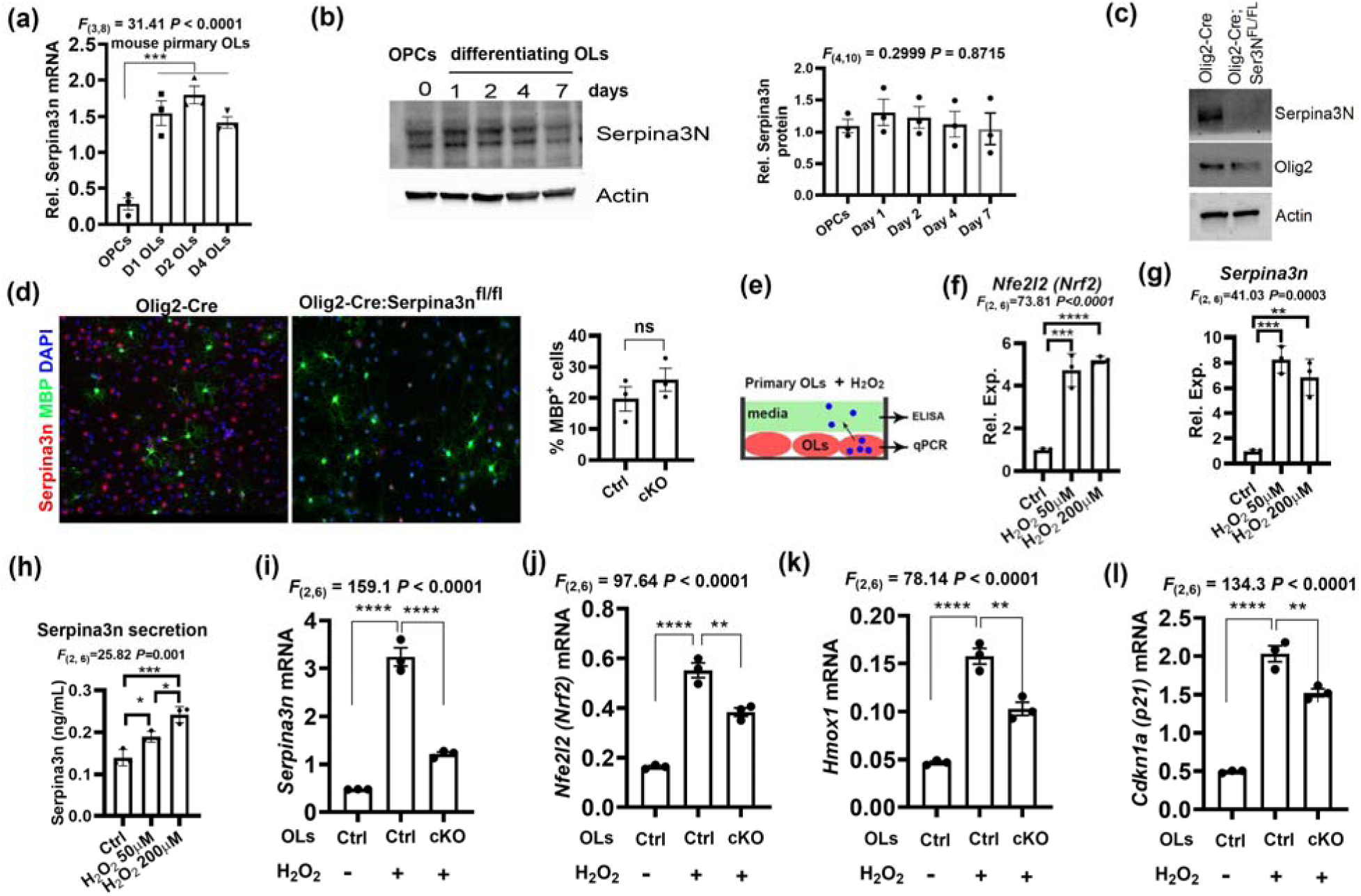
– oligodendrocytes enhance Serpina3n expression and secretion in response to oxidative stress *in vitro*. **(a)**, RT-qPCR assay for Serpina3n mRNA in primary mouse OPCs and differentiating OLs at day 1 (D1), D2, and D4 in differentiation media. **(b)**, Western blot assay for Serpina3n protein in primary rat OLs at different time points. **(c)**, Western blot assay for Serpina3n in D2 primary OLs from Serpina3n cKO and Ctrl brain. (d), double fluorescent immunocytochemistry of Serpina3n and MBP in D2 primary OLs and percentage of MBP^+^ OLs among DAPI^+^ cells. **(e)**, primary mouse OLs at D0 were treated with 50µM or 200 µM hydrogen peroxide H_2_O_2_ for 24 hours for downstream analyses. **(f)**, Serpina3n expression quantified by RT-qPCR. (**g**) ELISA assay for Serpina3n secretion in culture media. (**h**) response of oxidative stress marker Nfe2l2 (Nrf2) to H_2_O_2_ assessed by RT-qPCR. (i) RT-qPCR assay for Serpina3n in D1 primary mouse OLs from Serpina3n cKO or Ctrl brain that were treated with 50µM H_2_O_2_ for 24 hours. **(j)**, responses of oxidative stress marker Nfe2l2 (left) and its target gene Hmox1 (right) to 50µM H_2_O_2_ (24 hours). **(k)**, RT-qPCR assay for the expression of the cell senescence markers Cdkn1a (p21) in 50µM H_2_O_2_^-^treated Serpina3n cKO and Ctrl primary OLs (24 hours). One-way ANOVA followed by Tukey’s multiple comparison test, * P<0.05, ** P<0.01, *** P<0.001, **** P<0.0001.

Consistent with our *in vivo* observations, Serpina3n-deficient OLs differentiated normally as Serpina3n-sufficent OLs did (**Fig. 10d**), suggesting that Serpina3n plays a dispensable role for normal oligodendroglial development.

We next determined if Serpina3n plays a significant role in regulating the injury state of oligodendrocytes under pathological conditions. Oxidative stress is commonly observed in the CNS of various pathological diseases/injuries (Allen and Bayraktutan, 2009; Chen et al., 2012; Fesharaki-Zadeh, 2022; Liguori et al., 2018; Ljubisavljevic, 2016). We exposed primary OLs with hydrogen peroxide (H_2_O_2_) (**Fig. 10e**), a well-known inducer of cellular oxidative injury (Ransy et al., 2020). Oxidative injury to oligodendrocytes was verified by the marked induction of the oxidative stress-responsive transcription factor NRF2 (gene symbol *Nfe2l2*) (Ma, 2013) (**Fig. 10f**). We found that Serpina3n expression was remarkably upregulated in H_2_O_2_-treated OLs (**Fig. 10g**). ELISA assay showed that the level of Serpina3n in the culture media was significantly increased compared with the non-injured condition (**Fig. 10h**), which is in line with the secretory feature of Serpina3n. These data suggest that oligodendrocytes respond to oxidative injury by upregulating Serpina3n expression and secretion. Interestingly, the response of OLs to oxidative injury seemed to reach the plateau at 50µM H_2_O_2_ because no further increase in the expression of Serpina3n and Nrf2 was observed compared with 200µM H_2_O_2._ We therefore used 50µM H_2_O_2_ for our subsequent cKO experiments.

To demonstrate a role of Serpina3n in regulating oligodendrocyte injury, we cultured oligodendrocytes from Serpina3n cKO or Ctrl mice and treated them with or without 50µM H_2_O_2_ (**Fig. 10i**). Nrf2 expression was significantly rescued in cKO OLs treated with H_2_O_2_ compared with Ctrl OLs under the same culture conditions (**Fig. 10j**), suggesting that Serpina3n promotes H_2_O_2_-induced oxidative injury. The attenuated oxidative injury in cKO OLs was further corroborated by the reduced expression of Nrf2 target gene Hmox1 (**Fig. 10k**). Oxidative stress induces cell senescence (Faraonio, 2022). Consistently, we found that H_2_O_2_ treatment induces the expression of Cdkn1a (p21), a canonical marker of cell senescence (Gonzalez-Gualda et al., 2021), in cultured Ctrl OLs. Impressively, Serpina3n cKO decreased Cdkn1a expression in H_2_O_2_-treated OL culture (**Fig. 10l**). Collectively, these data suggest that Serpina3n aggravates oxidative stress and cell senescence under injury conditions.

## Discussion

Serpina3n/SERPINA3 is an acute phase glycoprotein that is secreted primarily by the liver to the bloodstream in response to system inflammation (Jain et al., 2011). SERPINA3 is elevated in the plasma and/or cerebrospinal fluid of people affected by cancers (de Mezer *et al*., 2023) and neurological diseases/injuries such as Alzheimer’s disease (DeKosky et al., 2003) and multiple sclerosis (Fissolo et al., 2021). The physiological role of the immune-related protein Serpina3n in regulating neural development and neuropathology remains underdefined partially due to lack of genetic animal tools. A recent study attempted to answer this question by using gain-of-function approaches (Zhao *et al*., 2022). It was reported that constitutive brain overexpression of human SERPINA3 increases neurogenesis and augments cognitive function in mice (Zhao *et al*., 2022). These important data suggest a crucial role of SERPINA3 in brain development and mouse behavior. In the current study, we found oligodendroglial lineage cells are the major cell population expressing Serpina3n in the postnatal brain. We leveraged the loss-of-function approaches to determine the physiological function of Serpina3n in brain development. Our results demonstrate that Serpina3n plays a minor role in postnatal brain cell development and animal motor/cognitive behaviors. The discrepancy of our data and those of the previous study (Zhao et al., 2022) suggest that Serpina3n may play a more important role in the development of embryonic neural precursor cells than postnatal brain cells. Alternatively, enforced overexpression of Serpina3n in cell types which otherwise lack Serpina3n expression, may account for the observed phenotypes (Zhao et al., 2022). Another possibility is that human SERPINA3 may have additional role to murine Serpina3n in brain development. Future studies are required to clarify these possibilities.

One of the significant findings of our current study is that Serpina3n protein was expressed primarily in oligodendroglia during early postnatal development of the murine CNS. This conclusion is proved by our Serpina3n cKO mice in which Serpina3n immunoreactive signals were abolished. Given the expression of Serpina3n in oligodendroglial lineage cells, we employed *Olig2-Cre:Serpina3n*^fl/fl^ transgenic mice to investigate its physiological role in in postnatal brain cell development (including oligodendroglial development) and animal behaviors. Serpina3n cKO mice did not show ataxia or tremor, which is the typical behavioral manifestation of CNS developmental hypomyelination. Our quantitative data showed that motor coordination, motor learning ability, regularity index of walking gait, and spatial learning and memory were all comparable in Serpina3n cKO to the control animals. Consistent with the normal behavior, our comprehensive analyses at the histological and molecular levels demonstrate overall comparable rates of oligodendrocyte differentiation and myelination in Serpina3n cKO mice to those in control mice. Based on these data, we conclude that Serpina3n plays a minor role, if any, in regulating oligodendroglial myelination and animal motor/cognitive function under homeostasis.

We hypothesize that oligodendroglia-derived Serpina3n regulates microglial and astroglial activity, which subsequently impacts brain neuronal development. However, our comprehensive analyses at the histological and molecular levels failed to identify impairment of microglial, astroglial, and neuronal development. These data suggest that the baseline expression of Serpina3n in oligodendrocytes plays a minor role in postnatal brain development. We further hypothesize that the baseline Serpina3n expression may prime oligodendrocytes for responding to CNS insults and regulate oligodendrocyte health under injured conditions. In support of this hypothesis, we found that oxidative stress remarkably increased Serpina3n expression and secretion in oligodendrocytes. Importantly, our cKO culture experiments point to a crucial role of Serpina3n in regulating the level of oxidative stress and cell senescence under injured circumstances. Our findings provide novel insights into the significance of Serpina3n in oligodendrocyte health under injury conditions. Given that Serpina3n has been dysregulated in various CNS pathologies (Zhu et al., 2024), it is intriguing to probe the potential roles of oligodendroglia-derived Serpina3n in regulating CNS pathophysiology. The genetic mouse lines generated in the current study provide powerful tools to study the *in vivo* functions of Serpina3n in CNS diseases/injuries.

## Data availability statement

The data generated or analyzed during this study are included in this article, or if absent, are available from the corresponding author upon reasonable request.

## Acknowledgements

We thank the funding agencies of NIH (R21NS125464, R01NS123080, R01NS123165, R01NS134887) and Shriners Hospitals for Children (85101-NCA-22, 85113-NCA-23).

## Declaration

The authors declared no conflict of interest and all consented to publication.

## Abbreviations

CNS, central nervous system; cKO, conditional knockout; OLs, oligodendrocytes; OPCs, oligodendrocyte progenitor cells; IHC, immunohistochemistry; MACS, magnetic-assisted cell sorting; LPS, lipopolysaccharide; CC, corpus callosum; SPC, spinal cord; EC, external capsule; CT, cortex; CB, cerebellum; i.p. intraperitoneal; SOX10, Sry-related HMg-Box gene 10; PDGFR- α: platelet-derived growth factor receptor A; Mag, myelin associated glycoprotein; Mobp, myelin associated oligodendrocyte binding protein; Mog, myelin oligodendrocyte glycoprotein; Cnp, 2’,3’-Cyclic nucleotide 3’-phosphodiesterase; Plp, proteolipid protein; MS, multiple sclerosis; P, postnatal day; PFA, paraformaldehyde; RT-qPCR, real-time quantitative PCR; TCF7l2, Transcription factor 7- like 2; TEM, transmission electron microscopy.

